# Mutations in *BBS2* Cause Apparent Nonsyndromic Retinitis Pigmentosa

**DOI:** 10.1101/2020.06.03.130518

**Authors:** Meghan DeBenedictis, Joseph Fogerty, Gayle Pauer, John Chiang, Stephanie A. Hagstrom, Elias I. Traboulsi, Brian D. Perkins

**Author notes:** <u>Corresponding author:</u> Brian D. Perkins, Ph.D., Department of Ophthalmic Research, Cleveland Clinic, Cleveland, OH 44195, USA.

## Abstract

**Purpose:** To identify and functionally test the causative mutations in the *BBS2* gene in a family presenting with retinitis pigmentosa and infertility and to generate a *bbs2^−/−^* mutant zebrafish.

**Methods:** A female proband and her male sibling were clinically evaluated and genetic testing with targeted next-generation sequencing was performed. Mutations were verified by Sanger sequencing. Protein localization was examined by transient expression and immunocytochemistry in cultured HEK-293T cells. Mutations in the zebrafish *bbs2* gene were generated by CRISPR/Cas9 and retinal phenotypes were examined by immunohistochemistry.

**Results:** The proband and her brother exhibited reduced visual fields, retinal degeneration, and bone spicule deposits, consistent with retinitis pigmentosa. The brother also reported symptoms consistent with infertility. Compound heterozygous mutations in the *BBS2* gene; namely NM_031885.4 (BBS2):c.823C>T (p.R275X) and NM_031885.4 (BBS2):c.401C>G (p.P134R), were identified in the proband and her brother. Both mutations interfered with ciliary localization of Bbs2 in cell culture. Mutation of the zebrafish *bbs2* gene resulted in progressive cone degeneration and rhodopsin mislocalization.

**Conclusion:** Missense mutations of *BBS2* leads to non-syndromic retinitis pigmentosa, but not Bardet-Biedl Syndrome, even though Bbs2 fails to localize to cilia. In zebrafish, the complete loss of *bbs2* results in cone degeneration and ciliopathy phenotypes, indicating a requirement for Bbs2 in photoreceptor survival.

## Introduction

Bardet-Biedl Syndrome (BBS) is a rare, autosomal recessive group of disorders that exhibits significant phenotypic and genetic heterogeneity. BBS results from dysfunction of the cilia and is referred to as a ciliopathy. The clinical features of different ciliopathies often overlap, making the diagnosis of BBS challenging in some cases. BBS is characterized by the primary features of postaxial polydactyly, truncal obesity, retinal degeneration, cognitive impairment, renal anomalies, and hypogonadism. Secondary features include among others hypertension, diabetes mellitus, hepatic fibrosis, hearing loss, dental abnormalities (hypodontia and dental crowding), congenital heart defects, brachydactyly, speech disorder, ataxia, and polydipsia-polyuria. A clinical diagnosis of BBS requires the presence of four of the primary features or three primary and two secondary features [1]. The prevalence of BBS differs between populations, with an incidence of approximately 1/160,000 in European populations [2] to as high at 1/13,000 among Bedouins in Kuwait [3].

Mutations in at least 21 different genes cause BBS and many of these genes encode proteins that regulate ciliary protein trafficking [4]. BBS2 (OMIM 606151) is one of eight highly conserved BBS proteins (BBS1, 2, 4, 5, 7, 8, 9, and 18) that form the BBSome, a protein complex that functions as a “ciliary gatekeeper” and regulates the entry and removal of specific membrane proteins from the cilium [5, 6]. The pathological features of BBS are thought to be linked to mislocalization and/or defects in ciliary trafficking of various proteins, including several G-protein coupled receptors (GPCRs), the leptin receptor, and the insulin receptor [7–12]. Nonsense, frameshift, and missense mutations in *BBS2* can result in BBS [13, 14].

The most penetrant feature of BBS is retinal dystrophy, with almost 97% of patients exhibiting some form of retinopathy [15]. Patients typically develop night blindness in the first decade of life and subsequently lose peripheral vision and visual acuity, all symptoms consistent with retinitis pigmentosa [4, 16]. Retinitis pigmentosa (RP) is a genetically heterogeneous condition with multiple inheritance patterns, including autosomal recessive (30-40% of cases), autosomal dominant (50-60%), and X-linked (5-15%) and is the most common hereditary retinal degenerative disease [17]. While the majority of RP cases remain isolated to the eye, up to 30% of patients can also exhibit extra-ocular manifestations indicating a syndromic association of the retinal dystrophy; this group of conditions includes BBS that accounts for up to 5% of all RP cases [17]. Here we describe a proband with compound heterozygous missense and nonsense variants in *BBS2* who presented with apparent nonsyndromic RP and her brother, who was affected by RP and infertility. These mutations disrupted ciliary localization of BBS2 protein in cultured cells. The generation of zebrafish with truncating mutations in *bbs2* resulted in progressive retinal degeneration and phenotypes consistent with cilia defects.

## Methods

### Clinical Evaluation

Retinal examinations, DNA sample acquisition, and molecular analysis were approved by the Cleveland Clinic Institutional Review Board (IRB). Written informed consent was obtained from the patients and was in accordance with HIPAA regulations and the tenets of the Declaration of Helsinki. The proband was evaluated in a specialized retinal dystrophy clinic. Family and medical histories were obtained and reviewed. Fundus photography, Goldmann visual field testing, and optical coherence tomography were performed.

### Molecular analysis

The proband’s DNA was sequenced for mutations in the retinal dystrophy genes via next generation sequencing (NGS) in a Clinical Laboratory Improvement Amendments (CLIA)-certified laboratory. Her brother’s and parent’s samples underwent targeted variant analysis in the Hagstrom laboratory. Genomic DNA was isolated from leukocyte nuclei prepared from blood samples as recommended by the manufacturer (QuickGene-610L; autogen). Saliva was collected in OG-250 Oragen DNA collection kits (DNA Genotek, Ottawa, ON, Canada) and processed using the PrepIT*L2P protocol (DNA Genotek, Ottawa, ON, Canada). Genetic analysis was performed with NGS via analysis of an inherited retinal dystrophy gene panel to identify the underlying cause of disease. 131 RP genes were known and screened at the respective time of analysis.

Following the identification of the mutation in the proband, targeted analysis was performed in the family’s samples. The DNA was amplified for exons 3 and 8 of the BBS2 gene (AF342736). Primer pairs were designed for both exons. (Exon 3 forward: CTGTTTTACTCAAAATCTGCTCAG and reverse: AAAGTAAAAATGCTTAAGGGTACC; Exon 8 forward: AGAATACTCTTGAAAACTGCTAGA and reverse: CTTTTTAAGGATTTTTCTCATCCC) Amplification was performed at an annealing temperature of 54 °C using Choice TM Taq Mastermix DNA Polymerase (Denville Scientific, Inc, Metuchen, NJ). PCR products were analyzed by direct sequencing using an automated sequencer. (3130XL, Life Technologies Corporation Grand Island, NY). In silico analysis of the identified missense variants was conducted using the Polyphen-2, pMut, and SIFT algorithms.

### Disease gene collections and targeted capture probes design

283 genes for 146 monogenic eye diseases were collected by systematic database (GeneReviews, OMIM, and RetNet) and literature searches, and expert reviews. Customized oligonucleotide probes were designed to capture the exons and 30 base pairs of adjacent sequences using NimbleGen (Roche) online oligonucleotide probe design system.

### Targeted sequencing library preparation and sequencing

Targeted sequencing libraries were prepared as following: 1 μg genomic DNA was sonicated to 200~300 bp sized fragments. This was followed by end-repair, A-tailing, Illumina adaptors ligation, and 4 cycles of pre-capture PCR amplification and sample indexing. The indexed PCR product of 20-30 samples were pooled, targeted capture was performed by hybridizing with capture probes, followed by 15 cycles of PCR amplification and validation of library products for sequencing. DNA sequencing was performed on Illumina HiSeq2000 sequencers to generate paired-end reads including 90 bps at each end and 8 bps of the index tag.

### Data filtering and analysis

Image analysis and base calling were performed using the Illumina Pipeline. Indexed primers were used for the data fidelity surveillance. Only reads that matched theoretical adapter indexed sequences and theoretical primer indexed sequences with no more than three mismatches were considered as valid reads. The human reference genome was obtained from the NCBI database, version hg19. Sequence alignment was performed using BWA (Burrows Wheeler Aligner) Multi-Vision software package (Li and Durbin, 2009). SNPs were called using SOAPsnp (Li et al., 2009) and Indels were identified using the GATK IndelGenotyper (http://www.broadinstitute.org/gsa/wiki/index.php/, The Genome Analysis Toolkit).

### Animal Maintenance

Wild-type zebrafish of an AB/Ekkwill hybrid strain were maintained and housed at approximately 28 °C on a 14/10 light/dark cycle using standard procedures [18] and with approval by the Institutional Animal Care and Use Committee (IACUC) at the Cleveland Clinic.

### CRISPR/Cas9 gene editing

Potential CRISPR target sites were identified in exon 4 of the zebrafish *bbs2* gene using ZiFiT (http://zifit.partners.org/ZiFiT/) [19] and gRNA synthesis was performed according to the protocol by Talbot and Amacher [20]. Briefly, oligonucleotides for gRNA synthesis (5’ TAGGAGGAAACTGTGCTCTTCA 3’ and 5’ AAACTGAAGAGCACAGTTTCCT 3’) were annealed and ligated into plasmid pDR274 (Addgene #44250, Watertown, MA). Purified plasmid DNA was digested with DraI (New England BioLabs, Beverly, MA) and used for in vitro transcription reaction to generate gRNA. Zebrafish embryos were injected at the 1-cell stage with a 1 nL solution of *bbs2* gRNA (200 ng/μL) and Cas9 protein (10 μM; New England BioLabs, Beverly, MA). Mutagenesis was confirmed by High Resolution Melt Analysis (HRMA) in injected F_0_ animals at 3 days post fertilization (dpf). Remaining F_0_ injected animals were raised to adulthood and outcrossed to wild-type fish and HRMA was performed on DNA from F_1_ progeny to screen for mutations. F_0_ founders producing a high degree of mutant F_1_ progeny were kept and individual *bbs2* alleles identified by sequencing.

### Preparation of wild-type and mutant BBS2 constructs

The full-length human *BBS2* clone in the pCMV6-entry vector was obtained commercially (Origene; Clone ID: RC204337). Using this as a template, the human *BBS2* ORF was PCR amplified with gene specific primers and sub-cloned into the pCDNA3.1V5/His TOPO TA vector (Invitrogen, Carlsbad, CA). The GeneArt Site-Directed Mutagenesis system (Invitrogen, Carlsbad, CA) was used to engineer two human *BBS2* mutant alleles (P134R and R275X). The *BBS2^P134R^* mutant allele was engineered using mutant primer pairs 5’-ACATTGGGAGACATTTCTTCCCgTCTTGCGATTATTGGTGGCAAT and 5’ ATTGCCACCAATAATCGCAAGAcGGGAAGAAATGTCTCCCAATGT-3; while the *BBS2^R275X^* mutant allele was engineered using mutant primer pairs 5’-GAAGGTTGATGCTCGAAGTGACtGAACTGGGGAGGTCATCTTTAA and 5’-TTAAAGATGACCTCCCCAGTTCaGTCACTTCGAGCATCAACCTTC-3’.

### Cell line authentication

HEK-293T cells were authenticated by American Type Tissue Collection (ATCC; Manassas, VA) using short tandem repeat (STR) analysis. Briefly, seventeen STR loci and a gender determining locus were amplified using PowerPlex 18D Kit (Promega; Madison, WI) and analyzed using GeneMapper ID-X v1.2 software (Applied Biosystems; Foster City, CA). Samples were a 100% match to a reference of human HEK-293T cells.

### Cell culture, transient transfection and immunofluorescence

HEK-293T cells were cultured in DMEM medium containing 10% FBS. For localization studies, cells were seeded in 6-well dishes with glass cover slips. At 60% confluence, cells were transfected with 3 μg of either wild-type (*BBS2*-WT-V5) or mutant *BBS2* constructs (*BBS2^P134R^-V5* and *BBS2^R275X^-V5*) using FuGENE HD (Roche, Branford, CT) according to the manufacturer’s instructions. Ciliogenesis was induced 24 h posttransfection by serum starvation using 0.5% FBS. After 24h, cell medium was removed from cells and washed twice with cold 1X PBS. Cells were fixed with a 4% PFA solution prepared in 1X PBS and subjected to indirect immunofluorescence. Cilia were detected using an anti α-acetylated tubulin antibody (Sigma, clone 6-11B-1), while the BBS2-V5 fusion proteins were detected using anti V5 antibody (Abcam, Cambridge, MA). Nuclei were stained using DAPI in the mounting medium. All mages were obtained using an epi-florescent microscope (Olympus BX60) equipped with a U-M61002 Tri-pass cube for DAPI/FITC/Texas Red. All images were obtained under a 100X oil immersion lens. Experiments were carried out in duplicate and approximately 100 cells, from 10-15 fields, were counted and scored per transfection.

### Immunohistochemistry

Immunohistochemistry was performed as previously described [21]. Briefly, zebrafish larvae were fixed in 4% paraformaldehyde, equilibrated in PBS + 30% sucrose, and embedded in Tissue Freezing Medium (Electron Microscopy Sciences, Hatfield, PA), prior to cryosectioning. Adult eyes were likewise fixed 2 hours in PBS + 4% paraformaldehyde and equilibrated as described [22]. Transverse sections including or adjacent to the optic nerve were incubating in blocking solution (PBS + 2% BSA, 5% goat serum, 0.1% Tween-20, 0.1% DMSO) for 1 hr. For cone inner segment analysis, the length of five Zpr-1+ cells were measured near the synapse at regular intervals across the retina. Rod outer segment length was measured similarly with zpr3 imaging. Cone outer segments and PCNA+ nuclei were counted across the entire retina. All experiments used between 3 and 6 animals per genotype.

## Results

### Two siblings from a non-consanguineous Caucasian family display features of RP and no other common signs of ciliopathies

A 40 year-old female presented for evaluation after an initial diagnosis of RP. Her medical history was significant for recurrent vocal cord carcinoma but otherwise non-contributory. She reported that her brother, three children, and parents were all healthy with no complaints of vision loss. A dilated fundus exam showed characteristic findings of RP, including optic nerve pallor, attenuated vessels, and the presence of bone spicule pigment deposits in the periphery (Fig. 1A). Her best-corrected visual acuity was 20/25 OU, but visual fields were significantly restricted (Fig. 1B). Subjectively, the patient did not appear overweight. The subject reported having no cardiac, renal, or neurological problems. A full panel analysis of 13 recessive retinitis pigmentosa genes available at the time was obtained and included *BEST1, CNGA1, CNGB1, CRB1, EYS, LRAT, NR2E3, NRL, PDE6A, PDE6B, RHO, RPE65*, and *USH2A*; genetic testing did not detect any variants. Later, genetic testing via next generation sequencing of additional retinal dystrophy genes detected 2 variants in *BBS2*, a pathogenic nonsense variant, c.823C>T leading to p.R275X, and a likely pathogenic missense variation, c.401C>G leading to p.P134R, which was previously identified in an Indian family with non-syndromic RP [23]. In silico analysis performed on the missense variant suggested it to be likely pathogenic (Polyphen score 0.957; PMut – pathological; SIFT score 0.08; potentially tolerated). Cosegregation analysis determined the variants to be in *trans*, but also determined the proband’s reportedly unaffected brother was also positive for both variants (Fig. 2)

**Figure 1.**
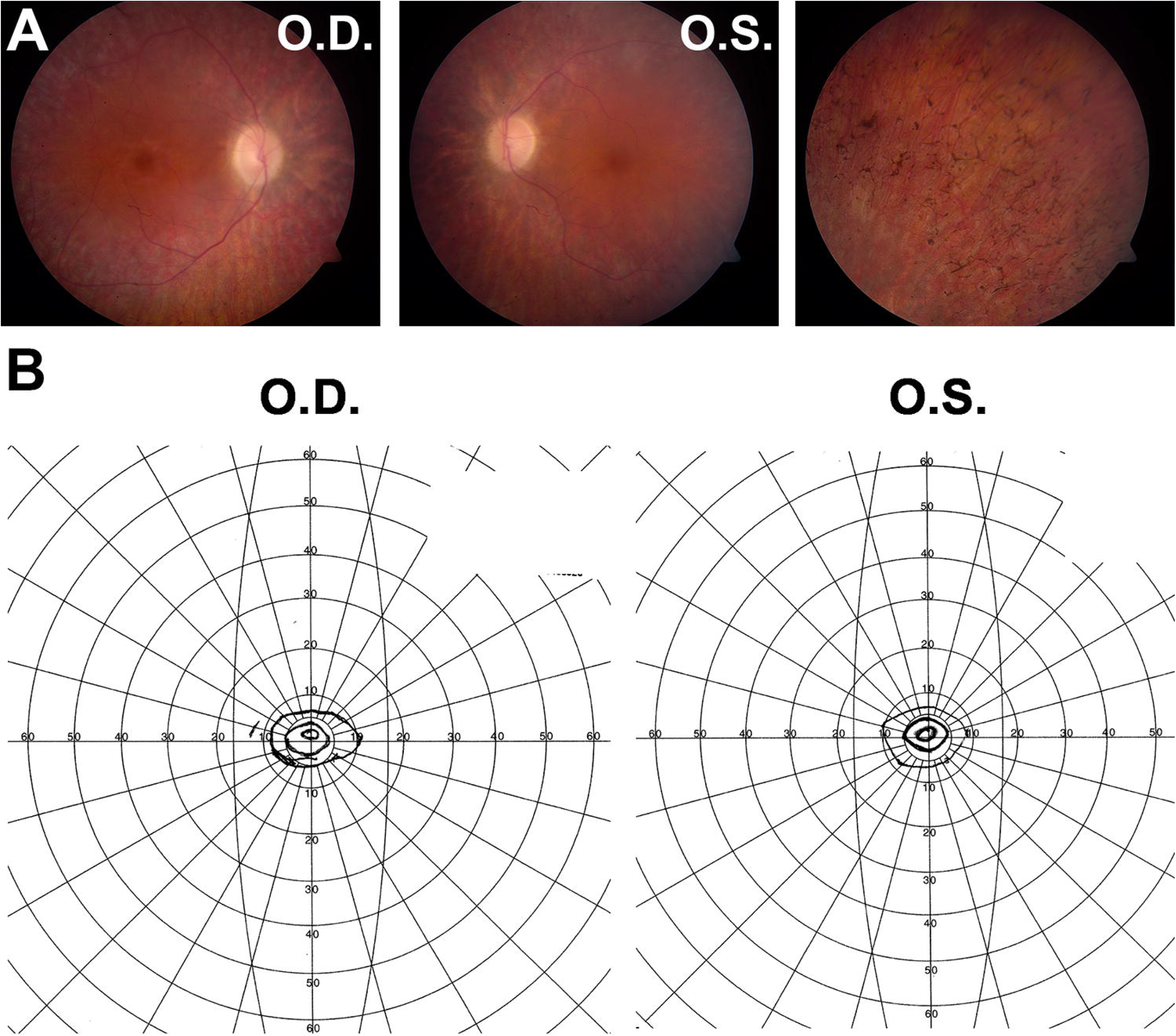
Evidence of retinitis pigmentosa and reduced visual fields in the proband (40 year old female). (A) Fundus photography showed optic disc pallor, vascular constriction and bone spicule deposits. (B) Goldmann visual field measurements of both eyes illustrate severely constricted visual fields.

**Figure 2.**
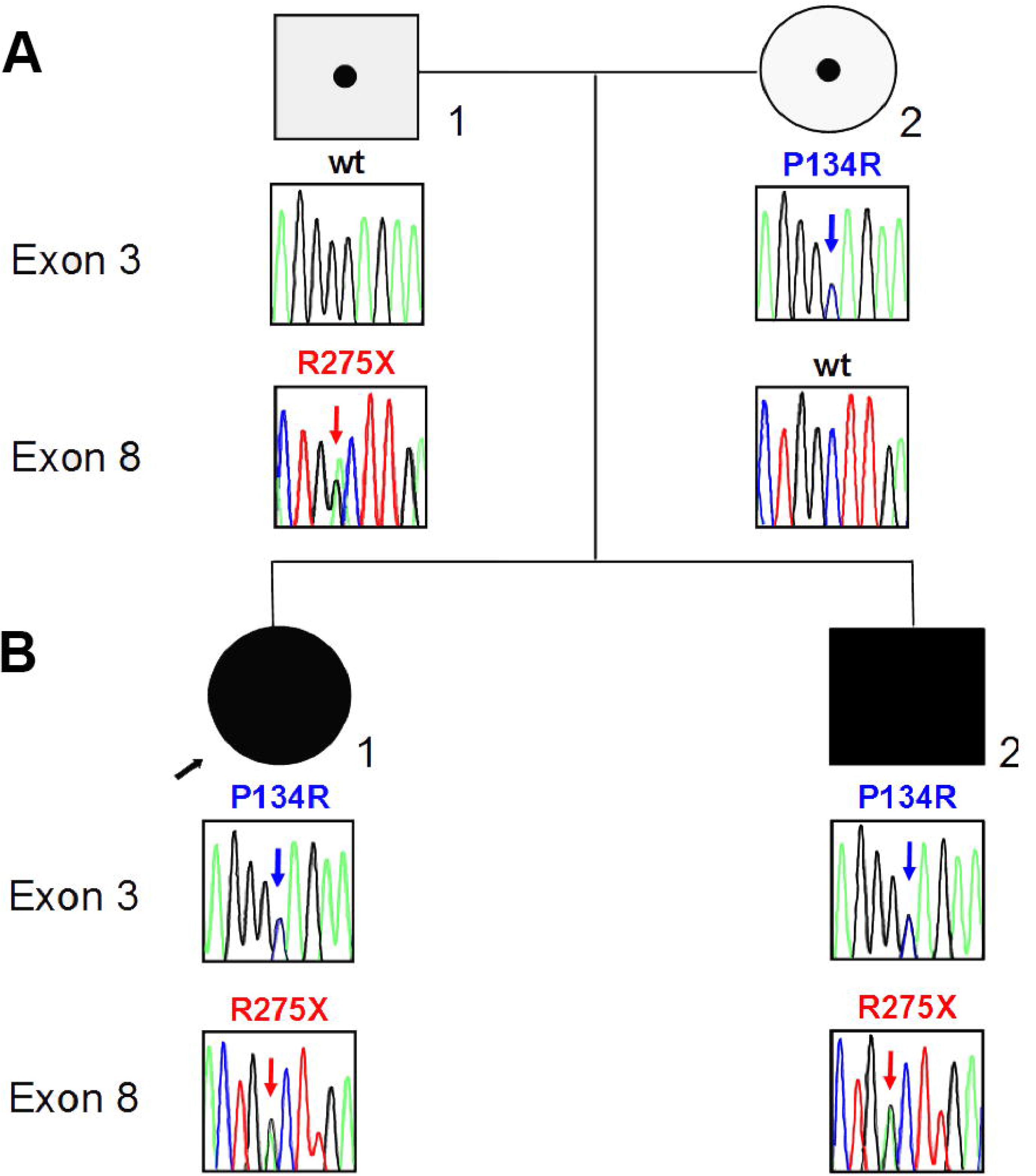
Pedigree of the proband, her brother and their parents. (A) Targeted variant analysis demonstrates the proband and her brother are both compound heterozygous for the p.P134R and p.R275X variants. (B) Parental testing confirms the variants are in *trans* with the mother and father carrying the p.P134R and p.R275X variants, respectively.

These results prompted us to revisit the family and perform a clinical evaluation on the brother, a 39-year old male. Although he did not report any complaints in visual function, dilated fundus examinations showed optic disc pallor, bone spicule pigment deposition, evidence of retinal degeneration, and mild peripheral visual field constriction (Fig. 3). During this examination he also disclosed fertility problems. His visual acuity was 20/20 OU. His confrontation visual fields were full and he had no color discrimination problems on Ishihara testing.

**Figure 3.**
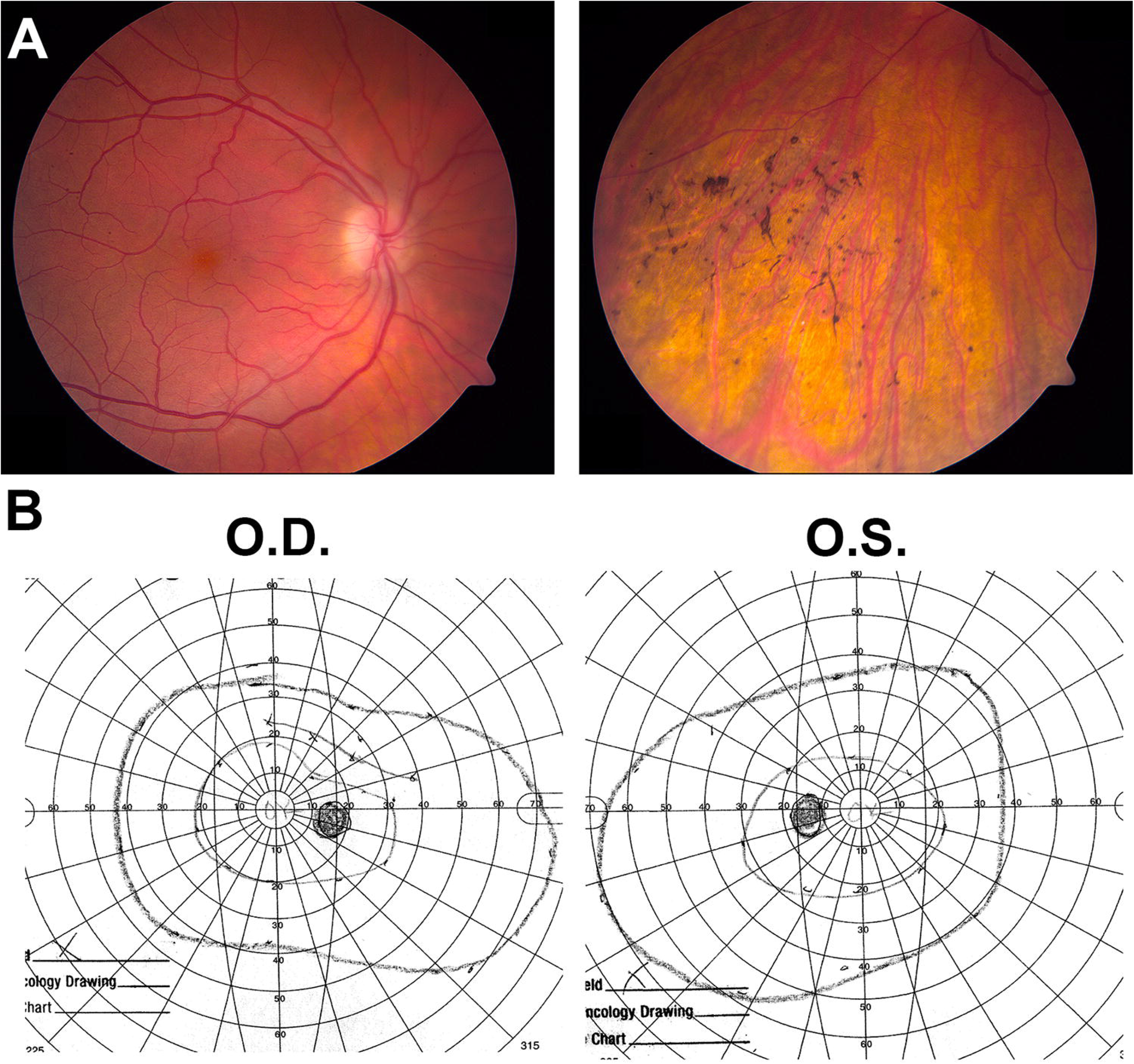
Evidence of retinitis pigmentosa and reduced visual fields in the proband’s brother, a 38 year old male. (A) Fundus photographs show a normal appearing optic nerve and posterior pole; however, pigmentary changes with bony spicules are seen in the inferior quadrants peripherally. (B) Goldmann visual field measurements of both eyes found only mildly reduced visual fields.

Additional testing was obtained following the molecular diagnosis of BBS to determine if the female patient had any systemic findings. Her free T4, TSH, and lipid panel were normal. Her complete metabolic panel and renal function panel were considered normal, although below the average reference range. Specifically, her fasting blood glucose was 62 mg/dL (reference range: 65 – 100 mg/dL) and her creatinine level was 0.58 mg/dL (reference range: 0.70 – 1.40 mg/dL). Prior imaging of her heart and reproductive organs did not reveal any abnormalities. Her most recent eye exam at the age of 48 years continued to show evidence of RP. Her best corrected visual acuity was 20/40 in both eyes and she was noted to have posterior subcapsular cataracts. Repeat metabolic testing was still normal.

The proband’s brother also had normal LH, FSH, Estradiol 17B, TSH and lipid panel levels. His blood pressure was normal. He had no history of recurrent pneumonia, bronchitis, or sinusitis. His complete metabolic panel was within normal limits. However, his testosterone levels were slightly below normal (173 ng/dL, reference range: 220 – 1000 ng/dL). Nigrosin-eosin staining of his sperm detected low amounts of viable sperm (40%, reference range: >75%). Complete semen analysis showed a decrease in motile sperm (30%, reference range: >40%) and a decrease in normal sperm heads (2%, reference range: 14-100%). The morphology of his sperm was also abnormal. Height was unavailable, but he weighed 270 pounds, placing him in the obese range. This patient was lost to subsequent follow up visits.

### The P134R and R275X mutations disrupts ciliary location BBS2 in HEK-293T cells

To investigate the functional consequences of the *BBS2^P134R^* and *BBS2^R275X^* alleles, plasmids encoding V5-tagged BBS2 proteins were introduced into HEK-293T cells (Fig. 4). The wild-type BBS2 protein was consistently seen localizing along the length of the cilium, as assessed by immunocytochemistry with antibodies against acetylated alpha tubulin and V5 (Fig. 4A). In contrast, the *BBS2^P134R^* mutant failed to localize to the cilia, was retained within the cytoplasm, and occasionally aggregated away from the cilium (Fig. 4B). The BBS2^*R275X*^ mutant also failed to localize to the cilia and additionally showed punctate staining near the cell surface (Fig. 4C). Taken together, these data suggested that these mutations interfered with proper targeting or localization of BBS2 to the cilium. Lack of ciliary localization may be responsible for the retinal phenotypes associated with mutated *BBS2* alleles.

**Figure 4.**
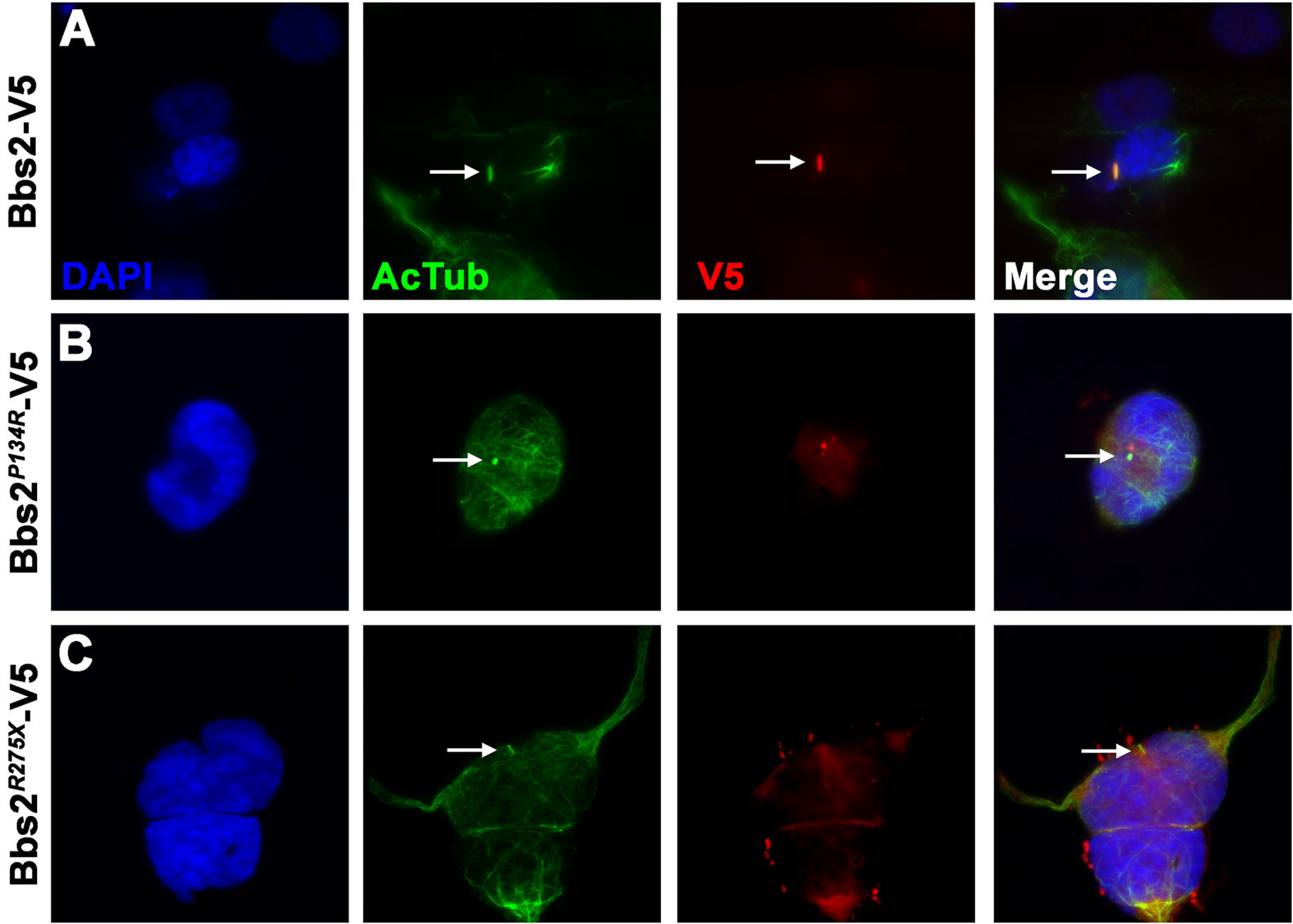
The P134R and R275X mutations disrupt ciliary location BBS2 in HEK-293T cells. Immunocytochemistry of HEK293-T cells expressing (A) wild-type *BBS-V5* (B) *BBS^P134R^-V5*, or (C) *BBS^R275X^-V5* mutant constructs using a monoclonal antibody again acetylated tubulin (AcTub – green) to label the ciliary axoneme and polyclonal antibodies against V5 (red) to label BBS-V5 fusion proteins. Nuclei are stained with DAPI (blue). Arrows note the location of the primary cilium.

### Mutation of zebrafish bbs2 results in progressive cone degeneration

To assess the role of *BBS2* in photoreceptor function, we utilized the CRISPR/Cas9 gene editing system to target exon 4 of the zebrafish *bbs2* gene. One mutant line (*lri82*) was recovered that resulted in the insertion of a 15-base pair fragment combined a 4-bp deletion, thereby yielding an 11-bp insertion and a frameshift with a premature stop codon (Fig. 5A). The truncated protein is predicted to contain the first 30% of the WD40 domain and then terminate after an additional frameshifted 58 amino acids (Fig. 5B). Founder (F_0_) animals were outcrossed to wild-type animals to generate *bbs2* heterozygote (*bbs2^+/−^*) zebrafish. Genotyping of clutches from *bbs2^+/−^* intercrosses revealed that homozygous offspring (*bbs2^−/−^*) appeared normal and were present in Mendelian ratios at 5 days post-fertilization (dpf) (Fig 5C). *bbs2^−/−^* mutants survived to at least 1 year of age, but developed spinal curvatures and were smaller than *bbs2^+/−^* and wild-type siblings (Fig 5D). These phenotypes resemble those of other zebrafish ciliopathy mutants [22].

**Figure 5.**
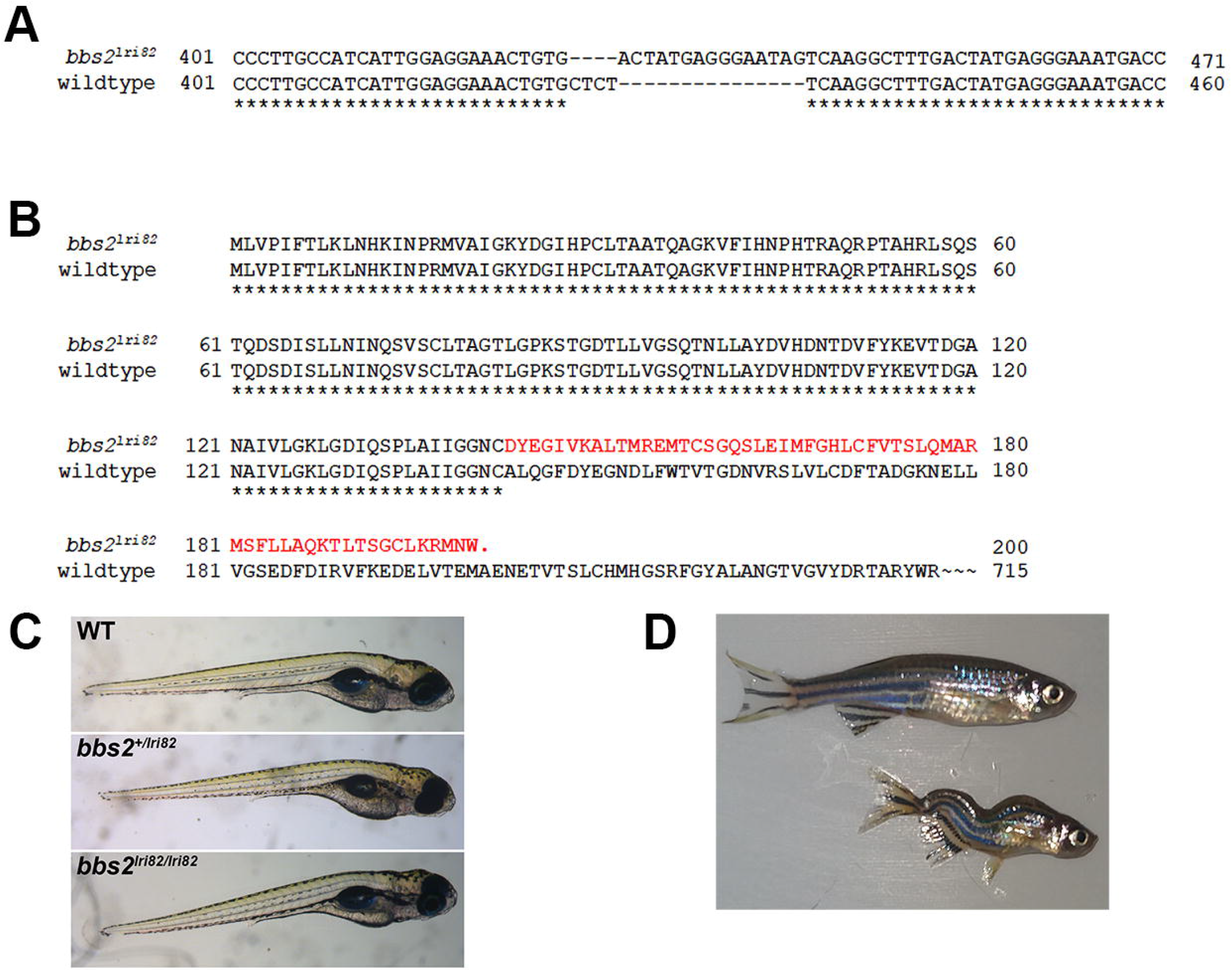
Mutation in *bbs2* exon 4 causes a frame shift and premature truncation. (A) Coding sequence alignment of the wild-type and *bbs2^lri82^* alleles. A 15bp fragment replaces “CTCT” from the wild-type allele, yielding a net 11-bp insertion and a frame shift. Numbering is relative to the start codon. (B) Predicted structure the of the wildtype and *bbs2^lri82^* peptides. The *bbs2^lri82^* mutant terminates after 58 frame-shifted codons (red). (C) Lateral view of 5 dpf wild-type (top), *bbs2^+/lri82^* heterozygotes (middle) and *bbs2^lri82/lri82^* homozygotes (bottom) that were verified by genotyping. No differences could be observed. (D) Lateral view of a 10 month old wild-type (top) and *bbs2^−/−^* homozygous mutant (bottom). Mutants are smaller and exhibit spinal curvature as adults.

The *bbs2^−/−^* zebrafish have normal retinal patterning at 5 dpf, but show early signs of photoreceptor dysfunction and long-term photoreceptor degeneration in adults. At larval stages (5 dpf), the inner segments of the red-green double cone photoreceptors were stained with the zpr-1 antibody, which recognizes arrestin 3-like [24], and with peanut agglutinin (PNA), which labels the extracellular matrix surrounding cone outer segments [25]. Although the number of cones and the size of the inner segments was no different between *bbs2^−/−^* mutants and wild-type siblings, the cone outer segments of *bbs2^−/−^* mutants were 20% shorter (5.43 ± 0.03 μm vs. 6.83 ± 0.18 μm, p=0.0015; Figs. 6A and 6B). Both juvenile *bbs2^−/−^* mutants (2.5 mpf) and adult *bbs2^−/−^* mutants showed signs of cone disorganization, with both cone inner segments and outer segments being significantly shorter than wild-type siblings (Figs. 6A, 6C, 6D) and reduced cone numbers in 6 mpf adults (Fig. 6D). In 6 mpf *bbs2^−/−^* mutants, rhodopsin was mislocalized to the rod inner segments, although the size of the rod outer segments was unchanged (Figs. 6E and 6F). This phenotype resembled that of the zebrafish *cep290^−/−^* mutant, which also showed signs of degeneration [22]. As retinal injury and cell death is known to stimulate regeneration of lost retinal neurons through the reprogramming and proliferation of Müller glia [26], retinal sections were stained with PCNA to identify proliferating cells. In wild-type retinas, only 1-2 PCNA+ cells were found in either the inner nuclear layer (INL), where the Müller glia reside, and 4-5 PCNA+ cells in the ONL that represent rod precursors [27]. In contrast, *bbs2^−/−^* zebrafish had a 4-fold increase in PCNA+ cells in the INL (10.8 ± 3.0 vs 2.5 ± 0.4) and a nearly 7-fold increase in PCNA+ cells in the ONL (35.8 ± 7.7 vs 4.8 ± 0.6; Figs. 6G and 6H). Increased proliferation of unipotent rod precursors in the ONL indicates regeneration of dying rod photoreceptors. The limited proliferation of Müller glia in the INL suggest that the slow degeneration in *bbs2^−/−^* mutants was not sufficient to trigger a robust regeneration response.

**Figure 6.**
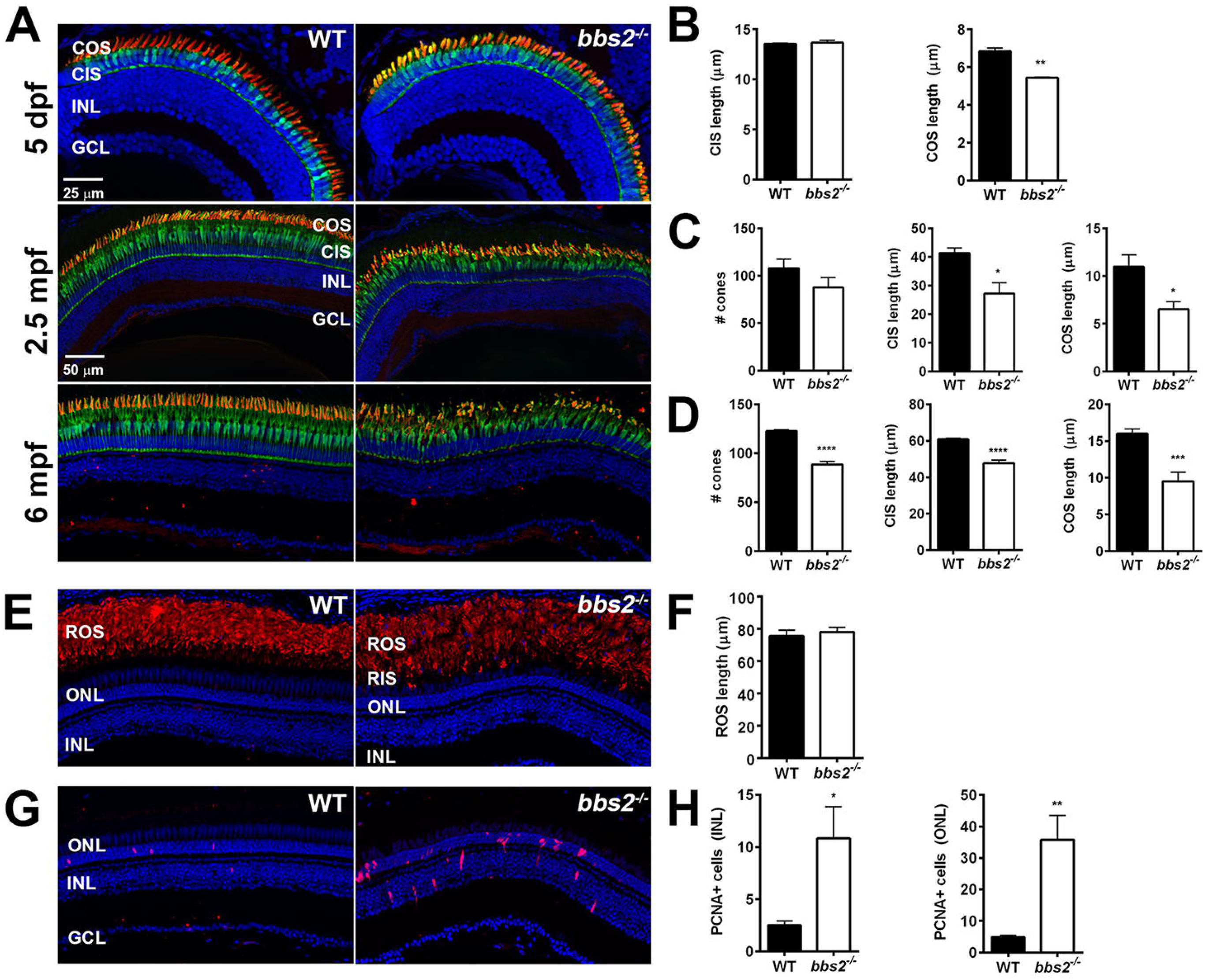
*bbs2^−/−^* zebrafish exhibit progressive cone degeneration. (A) Retinal sections were immunostained at the indicated time points with the monoclonal antibody zpr1 (green) to label cone inner segments (CIS), and with PNA (red) to label cone outer segments (COS). (B) Quantification of cone inner segment length and cone outer segment lengths at 5dpf, (C-D) Quantification of cone numbers and CIS and COS lengths in 2.5 mpf and 6 mpf adults. (E) Retinas from 6 mpf animals immunostained for zpr3 to label rod outer segments (ROS). (F) Quantified ROS length at 6 mpf. (G) Retinal cryosections from 6 mpf animals were immunostained for PCNA to label proliferating cells. (H) Quantification of PCNA+ cell counts at 6 mpf in the inner nuclear layer (INL) and outer nuclear layer (ONL). Asterisks indicate statistically significant differences, as determined by an unpaired t-test. *p<0.05; **p<0.01; ***p<0.001.

## Discussion

Herein, we report two siblings with compound heterozygous mutations in *BBS2* that result in apparently nonsyndromic retinitis pigmentosa, which on detailed investigation revealed associated male infertility and mild obesity in the male but no other syndromic signs of BBS in the proband or her brother. The two mutations detected in the present patients appear more likely to cause retinal dystrophy with little or no systemic findings. The P134R mutant allele was previously described in a non-consanguineous family with non-syndromic RP [23]. Although biallelic inheritance of the R275X allele was clinically diagnosed as BBS [14], the clinical manifestations were restricted to one or two systems and did not include obesity and polydactyly, two classic symptoms associated with BBS. The variability in expression of the genetic defect was not only in the systems involved, but also in the severity of the retinal degeneration. The female patient in the present report exhibited earlier onset of RP symptoms and more rapid progression compared to the male sibling.

Clinical and functional data from this study provides further evidence that pathogenic variants of the *BBS2* gene can present with apparent nonsyndromic RP or male infertility. It is likely that with advancing age the male patient would have shown symptoms of his retinal dystrophy. Nonsyndromic RP or RP with minimal systemic features has been described in patients with mutations in six of the BBS genes (*BBS1, BBS3/RP55/ARL6, BBS8/RP51/TTC8, BBS9, BBS12*, and *BBS14/CEP290*) [28–31]. The first report of this association came in 2008 when Cannon et al. reported brothers who were homozygous for the most common missense mutation in *BBS1*, p.M390R [31]. Brother 1 had RP, polydactyly, and some mild learning difficulties but he did not meet clinical diagnostic criteria for BBS [1]. Similarly, brother 2 had only RP and polydactyly. A second study that investigated the phenotypic spectrum of the common p.M390R *BBS1* variant found biallelic inheritance in two siblings with nonsyndromic RP [28]; one patient who was compound heterozygous for the p.M390R variant and a second missense pathogenic variant, p.L388P with nonsyndromic RP; three patients (3 homozygous and 1 compound heterozygous) were labeled as possibly nonsyndromic RP; and six patients (5 homozygous and 1 compound heterozygous) with mild BBS. There was only 1 patient with classic BBS in that study. A study of Saudi patients with RP found 4 patients in 1 family to be homozygous for the missense mutation, p.A89V in the *BBS3* gene [30]. All had RP without systemic findings. Four members of a family with a homozygous splice site mutation in *BBS8*, namely IVS1-2A>G had nonsyndromic RP [29]. Recently, Shevach et al. reported an association between missense pathogenic variants in the *BBS2* gene and nonsyndromic RP [23]. One of the missense variants they found is the same missense variant we detected in the present family, p.P134R. Unlike the patient in that report, the second pathogenic variant we report is a nonsense variant. This observation further broadens both the phenotypic and genetic spectrum of BBS alleles.

While BBS is largely an autosomal recessive disease, a number of reports have attributed phenotypic variability to triallelism [14]. In one study aimed at testing the hypothesis that triallelism causes clinical variability, a homozygous deletion of exon 6 in the *BBS9* gene was detected in three sisters [32]. One of these patients had all the primary features of BBS, while her two sisters had RP but none of the other BBS features. No other variants were found in any of the other BBS genes in this family. One study found a novel *BBS12* variant, p.S701X, in three affected individuals of a consanguineous family. All three had postaxial polydactyly and retinal degeneration. There were no renal or genital anomalies, obesity, or other symptoms consistent with a clinical diagnosis of BBS [33, 34]. Much has been published on the phenotypic heterogeneity in patients with *BBS14/CEP290* gene. This gene has also been reported to cause Leber Congenital Amaurosis [35], Senior Loken Syndrome [36], Joubert syndrome [36, 37], and McKusick-Kaufman syndrome [38]. Modifier alleles have been proposed to influence phenotypic severity of *CEP290*-associated disease [39], but mechanisms such as nonsense-mediated exon skipping may prove more predictive [40].

This also represents the first report of a loss-of-function phenotype for any *bbs* gene in zebrafish. Several prior studies utilized morpholinos to knock-down *bbs* gene function in *bbs1-14* and *bbs18* and reported defects in early embryogenesis [34, 41–48]. Unfortunately, interpretations of morpholino phenotypes, particularly those related to *bbs* function, have recently come under increased scrutiny due to potential off-target effects [49]. The phenotypes of the *bbs2^−/−^* mutant reported herein are consistent with other genetic mutants in zebrafish that affect cilia genes [22, 50, 51]. Importantly, the zebrafish *bbs2^−/−^* mutant undergoes progressive photoreceptor degeneration, consistent with mouse models [11] and humans. As zebrafish are capable of regenerating damaged photoreceptors following acute injury [52, 53], the *bbs2^−/−^* mutant may serve as a useful model for studies of regeneration in inherited retinal dystrophies.

In conclusion, we demonstrate that mutations in *BBS2* cause non-syndromic RP and infertility, that these mutations prevent ciliary localization of Bbs2 protein, and that zebrafish lacking *bbs2* function exhibit progressive photoreceptor degeneration. Although the phenotypes reported do not meet the diagnostic criteria for BBS, patients with RP and difficulties conceiving children from sperm problems should still be suspected of having BBS mutations. Advances in genetic testing technology, such as next generation sequencing, and broader diagnostic genetic testing panels have increased our ability to molecularly diagnose patients. Using more unbiased approaches for genetic diagnosis may lead to detection of variants in genes otherwise not considered as causative for a patient’s disease.

